# Increased adipose tissue indices of androgen catabolism and aromatization in women with metabolic dysfunction

**DOI:** 10.1101/2022.04.04.486987

**Authors:** Giada Ostinelli, Sofia Laforest, Denham Scott.G, Marie-Frederique Gauthier, Virginie Drolet-Labelle, Emma Scott, Frédéric-Simon Hould, Simon Marceau, Natalie.Z.M. Homer, Catherine Bégin, Ruth Andrew, André Tchernof

**Author notes:** ***Corresponding Author:*** Andre Tchernof, Ph.D., Quebec Heart and Lung Institute, School of Nutrition, Laval University, 2725 Chemin Sainte-Foy (Y-4212), Québec, QC, CANADA G1V 4G5.

## Abstract

**Background:** Body fat distribution is a risk factor for obesity-associated comorbidities, and adipose tissue dysfunction plays a role in this association. In humans, there is a sex difference in body fat distribution, and steroid hormones are known to regulate several cellular processes within adipose tissue. Our aim was to investigate if intra-adipose steroid concentration and expression or activity of steroidogenic enzymes were associated with features of adipose tissue dysfunction in individuals with severe obesity.

**Methods:** Samples from 40 bariatric candidates (31 women, 9 men) were included in the study. Visceral (VAT) and subcutaneous adipose tissue (SAT) were collected during surgery. Adipose tissue morphology was measured by a combination of histological staining and semi-automated quantification. Following extraction, intra-adipose and plasma steroid concentrations were determined by liquid chromatography, electrospray ionization tandem mass spectrometry (LC-ESI-MS/MS). Aromatase activity was estimated using product-over-substrate ratio, while AKR1C2 activity was measured directly by fluorogenic probe. Gene expression was measured by quantitative PCR.

**Results:** VAT aromatase activity was positively associated with VAT adipocyte hypertrophy (p-value_adj_ < 0.01) and negatively with plasma HDL-cholesterol (p-value_adj_ < 0.01), while SAT aromatase activity predicted dyslipidemia in women even after adjustment for waist circumference, age and hormonal contraceptive use. We additionally compared women with high and low visceral adiposity index (VAI) and found that VAT excess is characterized by adipose tissue dysfunction, increased androgen catabolism mirrored by increased AKR1C2 activity and higher aromatase expression and activity indices.

**Conclusion:** In women, increased androgen catabolism or aromatization is associated with visceral adiposity and adipose tissue dysfunction.

**DISCLOSURE SUMMARY:** AT obtained consulting fees form Bausch Health, Novo Nordisk and research funding from Johnson & Johnson Medical Companies as well as Medtronic and GI Windows for studies unrelated to this manuscript. The other authors have nothing to disclose.

## INTRODUCTION

Although obesity is a well-known risk factor for the development of a wide range of cardiometabolic alterations, research has demonstrated over the years that the distribution rather than total amount of adipose tissue, plays a determining role in the development of obesity-associated comorbidities. Excess deposition of adipose tissue within the mesentery and greater omentum, defined as visceral adipose tissue (VAT), has been identified as the most likely driver of cardiometabolic risk (1). Not only has VAT accumulation been linked to ectopic fat deposition (2,3), but its area rather than that of subcutaneous adipose tissue (SAT), is also associated with insulin resistance independently of body mass index (BMI), age, sex and ethnicity (4).

Recent work showed that adipose tissue dysfunction may mediate the association between body fat distribution and cardiometabolic health (4). Adipose tissue dysfunction may be defined as a set of molecular, cellular and morphological characteristics impeding the main function of the adipose organ, i.e to store excess energy (5). Among the features describing these alterations are adipocyte hypertrophy (6,7), reduced adipogenesis (8), decreased lipid storage (9,10), cellular senescence (11), pericellular fibrosis (12), altered adipokine secretion (13), and increased inflammation (14,15). During the last decade, genome-wide association studies found that genes related to adipose tissue functions such as adipogenesis and lipogenesis (16,17) play important roles in body fat distribution.

In humans, there are clear differences between the two sexes in terms of body fat distribution. Although women display more fat mass than men for any given waist circumference (WC), their lipids are preferentially stored in SAT rather than VAT (1,18). This difference is attenuated with age (18,19), leading to the hypothesis that estrogens and androgens may play a pivotal role in regulating some aspects of adipose tissue function. As reported in the literature, sex steroids are known to modulate some of the key processes such as lipolysis, lipogenesis, adipocyte differentiation and adipokine secretion (20,21). Androgens strongly impair adipogenesis and adipose commitment in primary human pre-adipocytes irrespective of the adipose depot and sex in individuals with (22) or without obesity (23). In addition, despite controversies found in the literature (22), testosterone treatment increases basal and stimulated lipolysis in abdominal SAT and VAT (24-26). On the other hand, estrogens hinder lipogenesis (27) by diminishing lipoprotein lipase activity (28) while modulating adipogenesis through increased preadipocyte proliferation (29) and SAT-specific adipogenesis (30) in women. Estrogens may also be implicated in energy partitioning, as they increase very-low-density-lipoprotein plasma clearance (31) and partially enhance muscle beta-oxidative capacity (32). Adipose tissue expresses several steroidogenic enzymes involved in androgen inactivation via the activities of aldo-keto reductase 1C2 (AKR1C2), their biosynthesis via AKR1C3, or estrogen biosynthesis by aromatase. Interestingly, mRNA expression of both *AKR1C2* and *CYP19A1* is stimulated by glucocorticoids (33,34), which have an additional role besides their implication in obesity and adipose tissue metabolism (35).

Our aim was to investigate if intra-adipose steroid concentrations and the expression or activity of steroidogenic enzymes associate with features of adipose tissue dysfunction in individuals with severe obesity. We hypothesized that androgen conversion is increased in participants with adipose tissue dysfunction.

## METHODS

### Participants

This study is a secondary analysis of a larger sample of participants that was used to examine food addiction in severe obesity (36,37). Briefly, bariatric surgery candidates with severe obesity (BMI ≥ 35 kg/m^2^) were recruited at the Quebec Heart and Lung Institute Research Center (CRIUCPQ). Patients were included if they were older than 18 years of age and met the criteria for elective bariatric surgery (either biliopancreatic diversion or sleeve gastrectomy). Patients with severe psychiatric conditions, those who were pregnant or lactating, had type 2 diabetes mellitus or underwent previous bariatric surgery were excluded. For the present study, we selected our sample of convenience by including participants who did not take corticosteroids and who also consented to take part into the Quebec Heart and Lung Institute Biobank and donate adipose tissue samples and plasma at the time of surgery. To mirror sex representation in bariatric candidates in North America (38), which is stable to an 80:20 women-to-men ratio, we included 31 women and 9 men in our study. Anthropometry, menopausal status, fasting plasma glucose and lipid profile, blood pressure as well as medication use (including birth control) were retrieved from medical records. We performed a priori sample size calculations based on published studies on adipose tissue morphology or features visceral obesity, with a focus on steroid enzyme gene expression and AKR1C2 activity. Power analyses on adipose tissue cells size and anthropometrics were completed after sample selection. Results indicated that a sample including between 6 and 35 individuals was sufficient to generate differences between subgroups or correlations with 80% power. This study was approved by the CRIUCPQ ethics committee (CER IUCPQ-UL number 2019-3218, 21758).

### Adipose tissue morphology

Adipose tissue was collected during surgery from the subcutaneous compartment and at the distal portion of the greater omentum and was immediately flash frozen in liquid nitrogen in the operating room. Frozen tissue was used for fixation in 10% buffered formalin at 4°C for 24h before paraffin inclusion (39). To assess adipocyte size, adipose tissue slides were stained with hematoxylin and eosin and photos were taken at 5x magnification. Adipocyte diameter was measured using a semi-automated program in ImageJ. The area of at least 100 adipocytes was measured in ImageJ (Rasband, W.S., ImageJ, U. S. National Institutes of Health, Bethesda, Maryland, USA, https://imagej.nih.gov/ij/, 1997-2018) and mean diameter was measured assuming that adipocytes were perfect circles (39).

To assess adipose tissue pericellular fibrosis, slides were stained with Picro-Sirius red (12). Threshold values were set per batch and calibrated using three internal standards of low (5.5%), medium (11.8%) and high (13.7%) mean percent pericellular fibrosis.

Adipose tissue function and visceral adiposity was determined using both the hypertriglyceridemic waist (HTGW) (40) and the age-specific visceral adiposity index (VAI) cut-offs for increased cardiometabolic risk (41). Briefly, individuals with HTGW were identified as having WC ≥ 90 cm and fasting triglycerides ≥ 2.0 mM (40), while VAI was calculated using the published, validated sex-specific equations (41).

### Quantification of steroids

For intra-adipose steroid quantifications, approximately 200 mg of adipose tissue was used, while plasma steroids were quantified in aliquots of 200 μL. Tissues and aliquots were stored at -80°C until the time of analysis. Glucocorticoids, estrogens and androgens in adipose or plasma were quantified using a ultra-high performance liquid chromatography (uHPLC) Shimadzu Nexera X2 system (UK) coupled to a Sciex QTRAP® 6500+ (SCIEX, Warrington, UK) with an electrospray ionization (ESI) interface.

Details on solid-phase extraction in adipose tissue can be found elsewhere (42). Briefly, frozen adipose tissue samples were enriched, together with standard solutions with internal radio-labelled standards. Following enrichment, adipose tissue was homogenized in ethanol:ethyl acetate (1:1). C18 SepPak columns (12 cc, 2 g; Waters, Wilmslow, UK) were used. Columns were washed with water and 5% methanol before methanol:acetonitrile (1:1) elution and then dried under flow nitrogen. To improve estrone and estradiol quantification in adipose tissue, estrogens were derivatized. The derivatization method has already been validated and published in both plasma and adipose (42). Following derivatization, samples were suspended in water:acetonitrile (1:1) and loaded to liquid chromatography vials. Lower limits of quantification can be found in supplementary Table 1 (**Table S1**). Accuracy, measured as percentage relative mean error, and precision, as percentage relative standard deviation, for estrogens and glucocorticoids have already been published (42), while for androgens, precision ranged between 14.8% and 6.1% and accuracy between 8.6% and 4.3%.

As far as supported liquid extraction and steroid quantification in plasma is concerned, our methods mostly rely on LC-MS/MS parameter already published (43). Briefly, isotopically labelled standards dissolved in methanol were added and thoroughly mixed to each sample. Samples were transferred to an automated sample processor (Biotage, Uppsala, Sweden), and 200 μL formic acid (0.1% v/v) was added. Metabolites were then eluted by the addition of dichloromethane/propan-1-ol (98:2) and the eluate was dried with oxygen-free nitrogen. The dry extracts were then reconstituted in 70:30 water/methanol solution, thoroughly mixed before being injected for LC-ESI-MS/MS analysis. Chromatographic separation was achieved on a Kinetex C18 column (3×150 mm, 2.6 μm; Phenomenex, UK) fitted with a KrudKatcher Ultra In-Line Filter (Phenomenex, UK) with water and methanol as mobile phase, with ammonium fluoride (50 μM) as modifier. Lower limits of quantification are found in **Table S1**. Precision ranged between 11.4% (5*α*-DHT) and 6.3% (estradiol) while accuracy between 5.8 (5*α*-DHT) and 7.0 (estrone).

### Cortisol Awakening Response (CAR)

CAR is an indirect measurement of the hypothalamic pituitary adrenal (HPA) axis activity and is defined as the rapid rise in salivary cortisol in the first 30-45 minutes following awakening (44). Participants were asked to sample saliva during the first 30 minutes following awakening time on three consecutive days. Samples were taken at the time of awakening (S1), 15 and 30 minutes after. Participants were asked not to drink, eat or brush their teeth before and during sampling time and to keep samples refrigerated until the next visit at the CRIUCPQ. Saliva was extracted from the cotton swabs by centrifugation and stored at -80°C until analysis. Salivary cortisol was measured by competitive enzyme-linked immunosorbent assay (ELISA) as per manufacturer instructions (Salimetrics, Carlsbad, CA USA; antibody ID: AB_2801306). Assay detection range was between 0.33nM to 82.77 nM and cross-reactivity with salivary cortisone was 0.13%.

### AKR1C2 activity

AKR1C2, also known as the 3*α*-hydroxysteroid type 3 (3*α*-HSD3), is a member of the aldo-keto reductase 1C family and mainly catalyzes the conversion of 5*α*-DHT into 3α-androstenediol, but also plays a small role in the catalysis of progesterone into 20α-hydroxyprogesterone (45). Intra-adipose AKR1C2 activity was indirectly measured by fluorescence emission of the chemical compound Cumberone (9-benzoyl-2,3,6,7-tetrahydro-1H,5H,11H-pyrano[2,3-f]pyrido[3,2,1-ij]quinolin-11-one), kindly synthetized by the Organic Synthesis Service (CHU de Québec research center, Québec, Canada). Briefly, tissues were homogenized in 6.67 μL/mg tissue potassium-phosphate buffer (100 mM K_2_HPO_4_, 100 mM KH_2_PO_4,_ pH = 6.0) as previously described (46). The fat-free portion of the homogenate was isolated by centrifugating the homogenate twice (20 627 x g, 16°C, 5 min). The reaction mixture (10 nM NADPH, 25 μM Cumberone) diluted in potassium-phosphate buffer, was added to the homogenate in order to reach a 1:11 dilution. Fluorescence emission by the probe was measured during a 12-hour kinetic experiment: at a 1.5-minute interval for the first 90 minutes, followed by a 30-minute and a 1-hour interval in the four-hour section of the kinetic experiment. Each sample was controlled for blood and NADPH content by subtracting values obtained in an individual-specific blank. AKR1C2 activity was calculated as the slope in the first 30% section of the reaction (46).

### Adipose tissue gene expression

To assess gene expression, approximately 100 mg frozen adipose tissue was homogenized in QIAzol buffer (Qiagen, Hilden, Germany) using Tissue Lyser (Qiagen, Hilden, Germany). RNA extraction was performed using RNeasy Lipid Tissue Mini Kit (Qiagen, Hilden, Germany) on RNeasy Mini Spin Columns with on-column DNase digestion (Qiagen, Hilden, Germany) to further purify RNA from possible DNA contamination. RNA concentration was determined using Biodrop (BioDrop Lrd., Cambringe, UK).

cDNA synthesis was performed on 195 ng RNA using iScript reverse transcriptase (Bio-Rad Laboratories, Hercules, CA, USA) as per manufacturer’s protocol. Finally, real-time quantitative polymerase chain reaction (RT-qPCR) was executed with 13 ng cDNA using SSO advanced SYBR Green supermix (Bio-Rad Laboratories, Hercules, CA, USA) with the following settings: denaturation (95°C, 20 s), annealing (58-65°C, 20 s), synthesis (72°C, 20 s) followed by fluorescence measurement (40 cycles). All experiments performed according to the Minimum Information for Publication of Quantitative Real-Time PCR Experiment (MIQE) guidelines (47).

**Table S2** lists the target, sequence and length of the primers used. Primers were designed using NCBI primer design tool (https://www.ncbi.nlm.nih.gov/tools/primer-blast/) and synthetized by IDT (Integrated DNA Technology, Coralville, IA, USA). Specificity and optimal annealing temperature for each primer pair were validated by gel electrophoresis combined with Cq and melt curve analysis.

### Statistical analyses

Adipose steroid concentrations values were expressed in pmol/kg tissue while plasma concentrations were expressed in pM (exception made for cortisol and cortisone values, which were expressed in nM). The product over substrate ratio was used as an estimate of enzymatic activity in adipose tissue. In particular, the cortisol:cortisone ratio was used as an indicator of the 11*β*-HSD1 oxoreductase activity, the estrone:androstenedione and estradiol:testosterone ratios as markers of aromatase activity, and the estradiol:estrone ratio as an estimate of 17*β*-HSD estrogenic activity. The cortisol awakening response (CAR) was calculated as the incremental area under the curve (AUCi) and the average value over the three sampling days was used (48).

Data were analyzed using R (48). Correlation plots were generated using the *corrplot* package version 1.1.25, while multivariate linear regression with *lme4* package version 0.84. Comparison between individuals with versus those without adipose tissue dysfunction was performed using either the Wilcoxon test or *t*-test. Features of adipose tissue dysfunction were compared using two-way ANOVA with post-hoc Tukey Honestly Significant Difference test.

## RESULTS

Our sample included 31 women and 9 men aged 39.5 ± 7.6 years and with a mean BMI of 50.1 ± 6.4 kg/m^2^. Most of the adipose tissue features measured in this study, such as steroidogenic enzyme activity estimates (49) and adipocyte cell size (6) are known to differ between the two sexes and for this reason, analyses of both sexes combined could not be performed. Unfortunately, due to some missing anthropometric data, analysis of adipose tissue dysfunction in men was not possible. Because of sample size, which would have given analyses limited meaning, data collected on male participants are presented in supplemental **Table S3**. Anthropometric and metabolic characteristics of female participants are listed in **Table 1**, while steroid hormone concentrations can be found in **Table 2**. They were all premenopausal, with a BMI of 50.6 ± 5.7 kg/m^2^ and a WC of 131.7 ± 11.2 cm. None of the women included in this study were diabetic or taking medications. However, 2 of them were diagnosed with polycystic ovary syndrome (PCOS) and 10 were taking anovulants. In women, an association between intra-adipose and plasma androgen, estrogen or glucocorticoid concentrations (data not shown) was not present.

**Table 1:** Anthropometric, metabolic and medication intake of the 31 women, analyzed according to visceral adipose index (VAI) age-specific cut-offs. All data are listed as mean ± standard deviation (SD) or, when stated, median (Mdn) ± interquartile rage (IQR).

|  | All Women (N=31) | Low VAI (N=13) | High VAI (N=18) | p-value |
| --- | --- | --- | --- | --- |
| Age (years) | 39.1 ± 7.0 | 40.4 ± 5.2 | 38.1 ± 8.1 | NS |
| BMI (kg/m<sup>2</sup>) | 50.6 ± 5.7 | 51.4 ± 6.3 | 50.1 ± 5.5 | NS |
| WC (cm) | 131.7 ± 11.2 | 129.5 ± 9.2 | 133.4 ± 12.4 | NS |
| Hip circumference (cm) | 148.2 ± 12.7 | 147.9 ± 14.2 | 148.4 ± 12.0 | NS |
| WHR | 0.89 ± 0.07 | 0.88 ± 0.08 | 0.90 ± 0.06 | NS |
| Neck circumference (cm) | 41.9 ± 3.2 | 41.8 ± 2.9 | 41.0 ± 3.5 | NS |
| SAT adipocyte cell diameter (μm) | 91.8 ± 12.3 | 90.0 ± 7.3 N=9 | 92.7 ± 14.3 N=17 | NS |
| VAT adipocyte cell diameter (μm) | 83.5 ± 14.1 | 75.2 ± 10.1 N=10 | 88.4 ± 14.1 N=17 | < 0.01 |
| SAT pericell. fibrosis (%) | 6.0 ± 1.7 | 6.1 ± 1.9 N=10 | 6.1 ± 1.7 N=17 | NS |
| VAT pericell. fibrosis (%) [Md±IQR] | 3.9 ± 1.7 | 4.6 ± 2.7 N=12 | 3.9 ± 1.7 N=17 | NS |
| Fasting glucose (mM) | 5.8 ± 0.6 | 5.9 ± 0.7 | 5.7 ± 0.6 | NS |
| HbA1c (%) [Mdn±IQR] | 5.7 ± 0.4 | 5.4 ± 0.4 | 5.7 ± 0.4 | NS |
| Fasting triglycerides (mM) [Md±IQR] | 1.55 ± 0.9 | 0.94 ± 0.2 | 1.86 ± 0.6 | < 0.001 |
| Total cholesterol (mM) | 4.66 ± 0.6 | 4.4 ± 0.6 | 4.8 ± 0.6 | NS |
| HDL-c (mM) | 1.2 ± 0.3 | 1.3 ± 0.3 | 1.1 ± 0.3 | NS |
| LDL-c (mM) | 2.75 ± 0.5 | 2.7 ± 0.5 | 2.8 ± 0.5 | NS |
| Systolic blood pressure (mmHg) | 133.2 ± 11.7 | 128.6 ± 10.1 | 136.5 ± 12.0 | NS |
| Diastolic blood pressure (mmHg) | 84.6 ± 8.5 | 81.5 ± 7.2 | 86.9 ± 8.8 | NS |
| Menopausal (Y:N) | (0:31) | (0:13) | (0:18) | NS |
| PCOS (Y:N) | (2:29) | (1:12) | (1:12) | NS |
| Taking hypoglycaemics drugs (Y:N) | (0:31) | (0:9) | (0:40) | NS |
| Taking hypolipidaemics drugs (Y:N) | (0:31) | (0:9) | (0:40) | NS |
| Corticosteroids (Y:N) | (0:31) | (0:9) | (0:40) | NS |
| Anovulants (Y:N) | (10:21) | (4:9) | (6:12) | NS |

Abbreviations: WC= waist circumference, WHR= waist-to-hip ratio, SAT= subcutaneous adipose tissue, VAT = visceral adipose tissue, HDL-c = high-density lipoprotein cholesterol, LDL-c = low-density lipoprotein cholesterol, PCOS= polycystic ovary syndrome, Md = median, IQR = inter-quartile range, Y = yes, N= no

|  | All women | Low VAI (N=13) | High VAI (N=18) | p-value |
| --- | --- | --- | --- | --- |
| Plasma cortisol (nM)<br>[Mdn ± IQR] | 217.7 ± 92.1 | 200.4 ± 64.6 | 220.0 ± 142.6 | NS |
| Plasma cortisone (nM) | 54.5 ± 19.3 | 51.7 ± 17.6 | 56.5 ± 20.6 | NS |
| Plasma androstenedione<br>(pM) |  | 2238.9 ± 958.4 | 2551.7 ± 867.5 | NS |
| Plasma testosterone (pM) | 751.4 ± 275.6 | 739.3 ± 311.4 | 761.2 ± 253.0<sup>i</sup> | NS |
| Plasma<br>dihydrotestosterone (pM) | 209.0 ± 82.2 | 204.1 ± 77.6 | 212.5 ± 87.4 | NS |
| Plasma estrone (pM) | 718.2 ± 471.7 | 647.2 ± 400.5 | 769.5 ± 522.3 | NS |
| Plasma estradiol (pM)<br>[Mdn ± IQR] | 432.5 ± 239.1 | 333.1 ± 111.0 | 407.2 ± 259.1<sup>j</sup> | NS |
| CAR AUCi<br>(nM*min)[Mdn ± IQR] | 92.4 ± 129.5 | 134.0 ± 147.7 | 68.7 ± 80.9 | NS |
| S1 mean (nM) [Mdn ±<br>IQR] | 11.5 ± 5.9 | 11.9 ± 4.6 | 11.0 ± 5.3 | NS |
| S2 mean (nM) [Mdn ±<br>IQR] | 14.3 ± 8.4 | 16.0 ± 8.6 | 12.7 ± 7.7 | NS |
| S3 mean (nM) | 16.0 ± 8.3 | 17.1 ± 7.1 | 13.1 ± 9.1 | NS |
| SAT cortisol (F)<br>(pmol/kg) [Mdn ± IQR] | 65591.0 ± 70957.0 | 25382.1 ± 80623.7<sup>a</sup> | 31689.4 ± 55364.6<sup>j</sup> | NS |
| SAT cortisone (E)<br>(pmol/kg) [Mdn ± IQR] | 23132.0 ± 23979.8 | 28070.3 ± 21393.6<sup>b</sup> | 7468.2 ± 24011.3<sup>i</sup> | NS |
| SAT<br>dehydroepiandrosterone<br>(pmol/kg) [Mdn ± IQR] | 75791.9 ± 144541.0 | 89827.5 ± 100838.8<sup>a</sup> | 67510.0 ± 135271.9<sup>i</sup> | NS |
| SAT androstenedione<br>(pmol/kg) [Mdn ± IQR] | 26592.0 ± 65735.8 | 42276.5 ± 66065.2 | 18232.6 ± 32877.6 | NS |
| SAT testosterone<br>(pmol/kg) [Mdn ± IQR] | 2545.0 ± 3317.4 | 1919.0 ± 3884.7<sup>c</sup> | 653.8 ± 1908.2 | NS |
| SAT dihydrotestosterone<br>(pmol/kg) [Mdn ± IQR] | 630.9 ± 450.8 | 630.9 ± 1616.6<sup>d</sup> | 642.3 ± 413.0<sup>g</sup> | N/A |
| SAT estrone (pmol/kg)<br>[Mdn ± IQR] | 5375.0 ± 3922.7 | 4653.4 ± 5978.8 | 3160.1 ± 6260.7 | NS |
| SAT estradiol (pmol/kg)<br>[Mdn ± IQR] | 3133.9 ± 2360.4 | 3801.7 ± 3147.5 | 1943.7 ± 1931.1<sup>i</sup> | NS |
| SAT F:E ratio | 5.1 ± 2.2 | 5.0 ± 2.3<sup>e</sup> | 5.1 ± 2.2<sup>k</sup> | NS |
| SAT A4:DHEA ratio<br>[Mdn ± IQR] | 0.24 ± 0.13 | 0.30 ± 0.10<sup>a</sup> | 0.23 ± 0.09<sup>k</sup> | NS |
| SAT E1:A4 ratio [Mdn ±<br>IQR] | 0.12 ± 0.10 | 0.10 ± 0.04 | 0.17 ± 0.11 | < 0.01 |
| SAT E2:T ratio [Mdn ±<br>IQR] | 1.6 ± 2.0 | 1.1 ± 1.1<sup>c</sup> | 2.5 ± 3.2<sup>i</sup> | < 0.05 |
| SAT (E1+E2) : (A4+T)<br>ratio [Mdn ± IQR] | 0.20 ± 0.11 | 0.16 ± 2.0<sup>c</sup> | 0.23 ± 4.4<sup>i</sup> | < 0.05 |
| SAT E2:E1 ratio [Mdn ±<br>IQR] | 0.6 ± 0.3 | 0.64 ± 0.30 | 0.59 ± 0.20<sup>i</sup> | NS |
| SAT AKR1C2 activity<br>(FU/min) | 13.5 ± 5.3 | 12.7 ± 5.8<sup>c</sup> | 14.0 ± 5.0 | NS |
| VAT cortisol (pmol/kg) | 64065.0 ± 57712.8 | 30469.2 ± 27671.6<sup>f</sup> | 67221.6 ± 85544.4 | NS |
| VAT cortisone (pmol/kg)<br>[Mdn ± IQR] | 23131.6 ± 30748.7 | 30061.8 ± 43544.8 | 11016.4 ± 21975.9 | NS |
| VAT<br>dehydroepiandrosterone<br>(pmol/kg) [Mdn ± IQR] | 108662.6 ± 112358.5 | 98305.6 ± 100605.4<sup>b</sup> | 119019.5 ± 115336.2<sup>k</sup> | NS |
| VAT androstenedione<br>(pmol/kg) [Mdn ± IQR] | 57212.0 ± 57337.6 | 45763.2 ± 115507.0 | 22328.4 ± 26708.3<sup>j</sup> | < 0.05 |
| VAT testosterone<br>(pmol/kg) [Mdn ± IQR] | 2045.0 ± 2068.1 | 1922.5 ± 3584.1<sup>c</sup> | 568.1 ± 2204.9 | < 0.05 |
| VAT dihydrotestosterone<br>(pmol/kg) [Mdn ± IQR] | 473.3 ± 547.7 | 472.4 ± 109.6<sup>g</sup> | 1137.1 ± 912.0<sup>d</sup> | N/A |
| VAT estrone (pmol/kg)<br>[Mdn ± IQR] | 5382.0 ± 3517.5 | 5338.9 ± 4595.7 | 3976.0 ± 6165.1 | NS |
| VAT estradiol (pmol/kg)<br>[Mdn ± IQR] | 3270.0 ± 2535.4 | 3067.6 ± 2461.1 <sup>c</sup> | 2364.8 ± 3008.1 <sup>i</sup> | NS |
| VAT F:E ratio [Mdn ± IQR] | 3.5 ± 2.5 | 3.8 ± 2.7 <sup>f</sup> | 3.4 ± 3.1 | NS |
| VAT A4:DHEA ratio<br>[Mdn ± IQR] | 0.23 ± 0.17 | 0.27 ± 0.22 <sup>b</sup> | 0.20 ± 0.14 <sup>k</sup> | NS |
| VAT E1:A4 ratio [Mdn ± IQR] | 0.10 ± 0.8 | 0.08 ± 0.3 | 0.13 ± 0.1 <sup>j</sup> | <0.01 |
| VAT E2:T ratio [Mdn ± IQR] | 2.0 ± 2.5 | 1.3 ± 1.9 <sup>h</sup> | 2.4 ± 3.4 <sup>i</sup> | NS |
| VAT (E1+E2) : (A4+T)<br>ratio [Mdn ± IQR] | 0.16 ± 0.11 | 0.11 ± 0.7 <sup>h</sup> | 0.20 ± 0.15 <sup>k</sup> | < 0.01 |
| VAT E2:E1 ratio | 0.57 ± 0.30 | 0.61 ± 0.33 <sup>c</sup> | 0.54 ± 0.28 <sup>k</sup> | NS |
| VAT AKR1C2 activity<br>(FU/min) | 8.3 ± 4.3 | 6.4 ± 3.6 <sup>h</sup> | 9.5 ± 4.4 <sup>j</sup> | < 0.05 |

Abbreviations: SAT= subcutaneous adipose tissue, VAT= visceral adipose tissue, VAI= visceral adiposity
index, CAR= cortisol awakening response, AUCi=incremental area under the curve, S1= first sample F=
cortisol, E= cortisone, DHEA= dehydroepiandrosterone, A4=androstenedione, T=testosterone, E1= estrone,
E2=estradiol, Md= median, IQR =interquartile range, FU= fluorescence units, min=minutes, NS= not significant,

First, we examined features of adipose tissue dysfunction. We assessed two morphological parameters already described by us and others as being associated with adipose tissue dysfunction: adipocyte size (6) and pericellular fibrosis (12). In addition, we measured mRNA abundance of genes encoding proteins involved in regulating adipose tissue function regulation: diacycl-glycerol acyl-transferase 2 (*DGAT2*) and glutathione peroxidase x3 (*GPX3*), which are respectively involved in triglyceride biosynthesis and adipose-specific insulin resistance, respectively. When comparing features of adipose tissue dysfunction in the two fat depots, we found that adipocytes were larger (91.8 ± 12.2 vs 83.5 ± 14.1 μm, p < 0.05) (**Figure 1A**) and that there was more pericellular fibrosis (6.0 ± 2.4 vs 4.0 ± 2.0 μm, p < 0.001) (**Figure 1B**) in SAT compared to VAT. Gene expression analysis (**Figure 1C**) revealed that *DGAT2* and *GPX3* expression were not significantly different in the two depots (p_anova_ > 0.05 for both). Because adipose tissue dysfunction has been proposed as a mediator of the link between abdominal obesity and cardiometabolic risk (50), we looked at associations between dysfunction features and cardiometabolic health. As illustrated in **Figure 1D**, we found that WC was positively associated with cell size in both SAT (r = 0.4, p < 0.05) and VAT (r = 0.5, p < 0.05). Much like VAT adipocyte diameter, WC was negatively associated with plasma HDL-c concentrations (r = -0.5, p < 0.01). Interestingly, *DGAT2* expression was not associated to any anthropometric or metabolic parameter, while *GPX3* expression was associated with a more favorable plasma lipid profile (**Figure 1D**).

**Figure 1:**
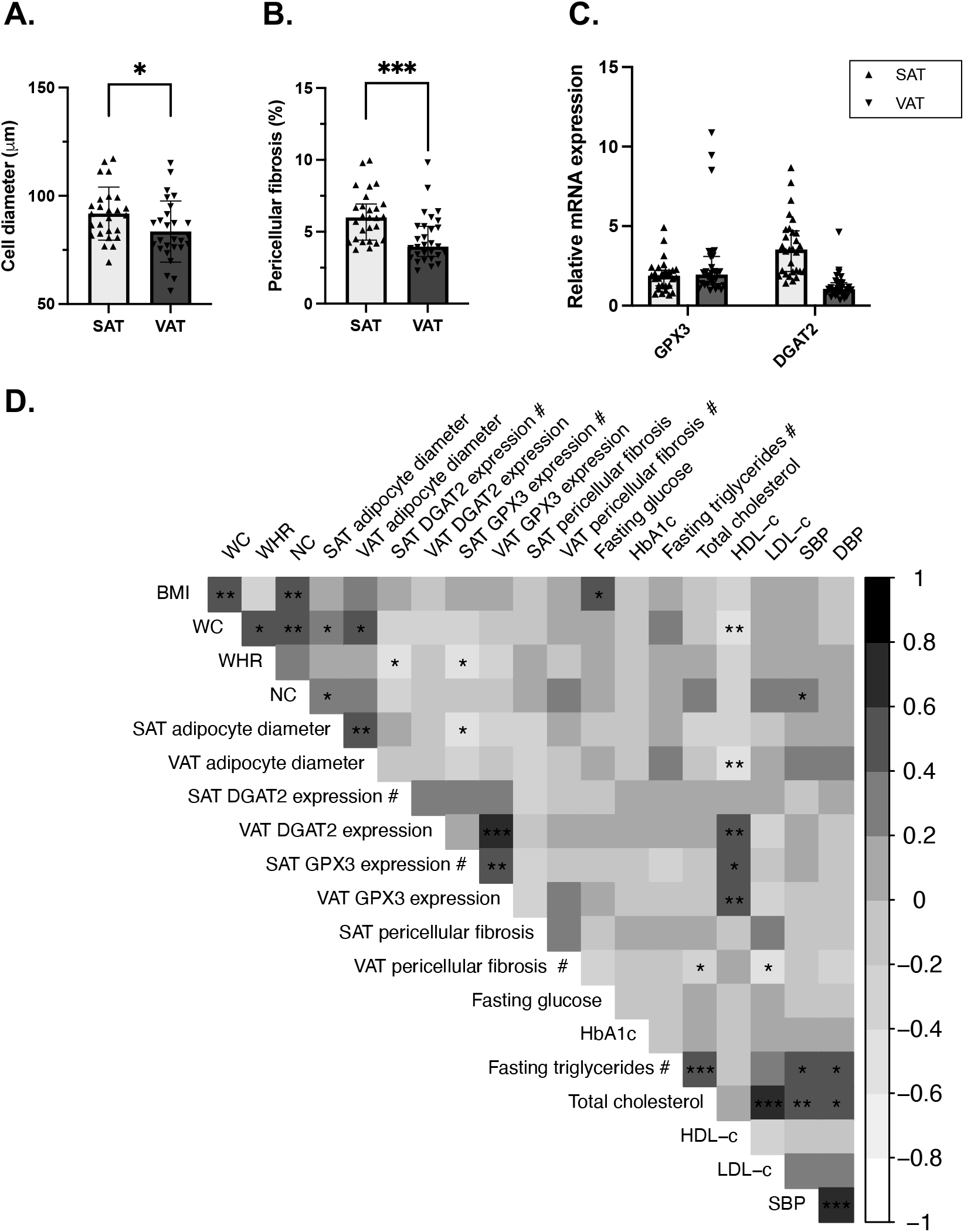
Features of adipose tissue dysfunction. Differences in mean adipocyte diameter (μm) (N=26, SAT vs. N= 27, VAT) (**A**), pericellular fibrosis (%) (N=27, SAT vs. N= 29, VAT) (**B**) and the expression of *DGAT2* (N=30, SAT vs. N= 30, VAT), implicated in triglyceride biosynthesis, and *GPX3* (N=30, SAT vs. N= 30, VAT), associated with lower intracellular reactive-oxygen species (ROS) production (**C**) in subcutaneous (SAT) and visceral adipose tissue (VAT). Pearson correlation coefficient range and p-values (**D**) for untransformed and log10-transformed (#) variables. Correlations with HbA1c were assessed with Spearman’s rank coefficients. The grey intensity reflects the strength of the correlation (Pearson’s r or Spearman’s rho), strong positive associations are illustrated in black while strong negative association in white.

Univariate analyses were conducted to uncover correlations between anthropometric variables and the activity or expression of steroidogenic enzymes. BMI was negatively associated with the estradiol-to-testosterone ratio in VAT (r = -0.4, p < 0.05) and with the estradiol-to-estrone ratio in SAT, an indirect measurement of 17*β*-hydroxysteroid dehydrogenases (r = -0.4, p < 0.05). Similarly, WC was also negatively associated with estradiol-to-estrone ratio in SAT (r = -0.4, p < 0.05) and AKR1C2 expression in SAT (r = -0.4, p < 0.05), while waist-to-hip ratio (WHR) was associated with the estrone-to-androstenedione ratio and the estrone + estradiol over testosterone + androstenedione ratio in SAT, both indicators of aromatase enzymatic activity (r = -0.4, p < 0.05 for both). In addition, neck circumference was negatively associated with SAT AKR1C2 expression (r = -0.4, p < 0.05). Finally, VAT adipocyte diameter was positively associated with the estrone-to-androstenedione ratio in VAT (r = 0.5, p = 0.01). No other significant association between anthropometric variables and steroidogenic enzymatic activities or expression were found.

We then tried to predict features of adipose tissue dysfunction using either plasma or intra-adipose steroid concentrations or steroidogenic enzyme activities (**Table 3**) in multiple linear regression models. Because the inclusion or exclusion of the two participants with PCOS did not alter the model, the analyses presented in **Table 3** include all 31 participants of the study.

**Table 3:** Steroid hormone concentrations in women predicts morphological features of adipose tissue dysfunction. Cortisone and intra-adipose estrogen are associated with adipocyte cell size and pericellular fibrosis. Intra-adipose estrone : androstenedione (E1:A4) ratio in women predicts features of adipose tissue dysfunction. Adjusted model takes into consideration waist circumference, age and anovulant drug use.

| Independent variable | Dependent variable | Unadjusted model |  |  |  |  | Adjusted model |  |  |  |  |
| --- | --- | --- | --- | --- | --- | --- | --- | --- | --- | --- | --- |
|  |  | Beta | SE | p-value | Adjusted R<sup>2</sup> | AIC | Beta | SE | p-value | Adjusted R<sup>2</sup> | AIC |
| Plasma cortisone | SAT adipocyte cell diameter | 0.31 | 0.11 | 0.01 | 0.20 | 202.30 | 0.31 | 0.11 | 0.009 | 0.29 | 201.52 |
| SAT cortisone | SAT pericellular fibrosis | 4.0E<sup>-05</sup> | 1.2 E<sup>-05</sup> | 0.002 | 0.34 | 79.34 | 4.1 E<sup>-05</sup> | 1.3 E<sup>-05</sup> | 0.005 | 0.24 | 84.98 |
| SAT estradiol | SAT pericellular fibrosis | 4.9 E<sup>-04</sup> | 1.1 E<sup>-05</sup> | 0.0002 | 0.44 | 88.86 | 5.6 E<sup>-04</sup> | 1.4E<sup>-04</sup> | 0.0006 | 0.42 | 92.41 |
| SAT estrone | SAT pericellular fibrosis | 2.5 E<sup>-04</sup> | 7.9 E<sup>-05</sup> | 0.001 | 0.31 | 100.37 | 2.7 E<sup>-04</sup> | 7.8 E<sup>-05</sup> | 0.002 | 0.29 | 103.37 |
| VAT E1: A4 | VAT adipocyte diameter | 84.97 | 32.38 | 0.01 | 0.19 | 210.67 | 89.80 | 32.12 | 0.01 | 0.35 | 207.10 |
| VAT E1: A4 | HDL-c | -1.86 | 0.71 | 0.01 | 0.16 | 14.84 | -1.94 | 0.64 | 0.005 | 0.44 | 5.10 |
| SAT E1: A4 | Triglycerides | 3.95 | 1.3 | 0.005 | 0.21 | 62.66 | 3.54 | 1.46 | 0.02 | 0.16 | 67.29 |
| SAT E1: A4 | HDL-c | -2.06 | 0.54 | 0.0006 | 0.31 | 8.29 | -1.53 | 0.55 | 0.009 | 0.40 | 6.25 |

Abbreviations: SE = standard error, AIC = Akaike information criterion, SAT = subcutaneous adipose tissue, VAT = visceral adipose tissue, HDL-c = high-density lipoprotein cholesterol

Our analysis revealed that, with an exception made for plasma cortisone concentrations, plasma steroid concentrations were not associated with any feature of adipose tissue dysfunction or metabolic impairment. On the other hand, VAT aromatase activity was associated with VAT hypertrophy, even after correction for WC, age and anovulant use (β_adjusted_ = 89.80, p_adjusted =_ 0.01). Similarly, aromatase activity in SAT was associated with dyslipidemia in our cohort, as demonstrated by the positive association with fasting plasma triglycerides and negative association with plasma HDL-c concentrations (p_adjusted_ < 0.05 and < 0.01 respectively). Generally similar results were found when considering aromatase activity as the sum of estradiol and estrone concentrations over the sum of testosterone and androstenedione. Finally, when looking at intra-adipose steroid concentrations, we found that SAT estradiol, estrone and cortisone concentrations were associated with SAT pericellular fibrosis.

To better capture any alteration occurring in adipose tissue dysfunction, we first divided the cohort using the HTGW algorithm (40). Neither adipocyte hypertrophy nor any of the measured markers of dysfunction were significantly different in participants who had versus those who did not have HTGW (data not shown). The VAI was used as another marker of adipose tissue dysfunction using age-specific cut-off values to identify individuals with increased cardiometabolic risk (41). We compared women having “High VAI”, N= 18 to those with “Low VAI”, N=13 (41) and found a significant difference in VAT adipocyte diameter (**Figure 2C**) but not in SAT (**Figure 2B**) or WC (**Figure 2A**). We found no difference either in SAT nor in VAT pericellular fibrosis (data not shown) and *DGAT2* expression (**Figure 2M**). However, women with adipose tissue dysfunction were characterized by lower expression in *GPX3* in both adipose tissue depots (**Figure 2M**). When looking at intra-adipose androgen concentrations, we found that women with adipose tissue dysfunction had lower VAT concentrations of both testosterone and androstenedione (**Figure 2D, E**). Interestingly, androgen aromatization was higher with adipose tissue dysfunction as illustrated by significantly increased aromatase activity ratios in SAT (**Figure 2F, G**), VAT (**Figure 2H**) and *CYP19A1* expression in VAT only (**Figure 2M**). The same difference in SAT and VAT was observed by inferring aromatase activity as the ratio of estrone + estradiol over testosterone + androstenedione (**Table 2**). Similarly, we observed increased 5*α*-DHT catabolism, as mirrored by an increased AKR1C2 activity (**Figure 2J**). This difference was not reflected in *AKR1C2* expression (**Figure 2M**). Increased aromatization of intra-adipose androgens was reflected by a trend for increased plasma estradiol concentration in women with high VAI (Md ± IQR: 333.15 ± 111.04 vs. 407.16 ± 259.10, p = 0.09). No difference was observed in intra-adipose estrone levels. Finally, consistent with the notion that adipose tissue dysfunction increases the risk of cardiometabolic diseases, we found that women with high VAI had atherogenic dyslipidemia, as illustrated by higher plasma Apolipoprotein B (ApoB) concentrations in **Figure 2L**. No differences were seen in terms of plasma and intra-adipose glucocorticoid concentrations, *HSD11B1* gene expression or in CAR between the two groups (data not shown).

**Figure 2:**
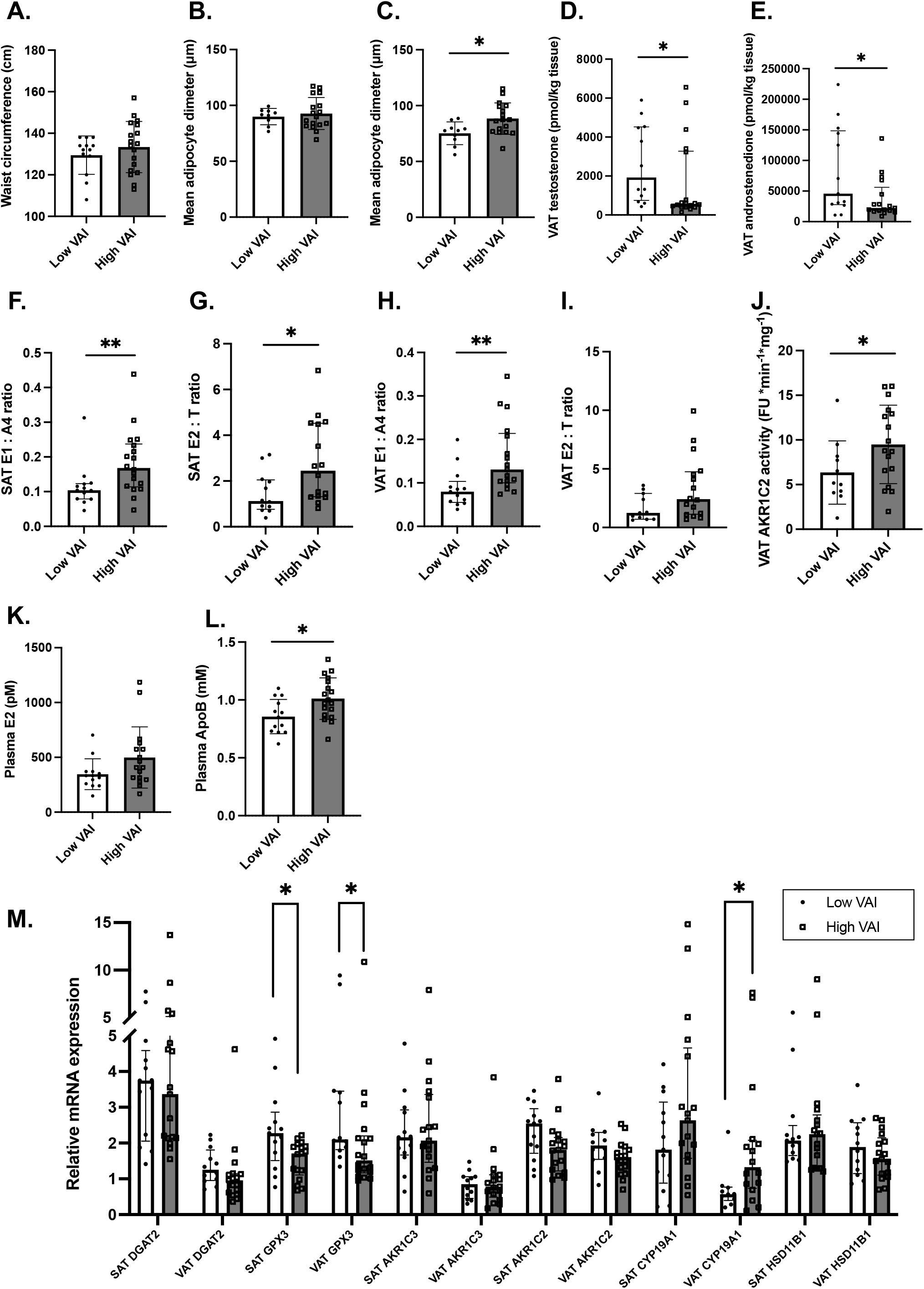
Comparison between women with high vs low visceral adipose index (VAI) in terms of adipose tissue dysfunction markers, steroid hormone concentrations and mRNA expression measurements. Differences waist circumference (A), SAT (low VAI, N= 9 vs. high VAI, N= 17) (B) and VAT (low VAI, N= 10 vs. high VAI, N= 17) (C) mean adipocyte diameter (μm), VAT testosterone (low VAI, N= 12) (D), androstenedione (high VAI, N= 17) (E) concentrations and comparison with aromatase activity indices (F), (low VAI, N= 12 vs. high VAI, N= 16) (G), (high VAI, N= 17) (H), (low VAI, N= 11 vs. high VAI, N= 16) (I), and AKR1C2 (low VAI, N= 11 vs. high VAI, N= 17) enzymatic activity (**J**). Differences in plasma estradiol (high VAI, N= 17) (K), and apolipoprotein B (L) as well as intra-adipose gene expression of *DGAT2* (SAT: high VAI, N=17; VAT: low VAI, N= 12), *GPX3* (SAT: high VAI, N=17; VAT: low VAI, N= 12), *AKR1C3* (SAT: high VAI, N=17; VAT: low VAI, N= 12 vs. high VAI, N=17), *AKR1C2* (SAT: high VAI, N=17; VAT: low VAI, N= 12 vs. low VAI =17), *CYP19A1*(SAT: high VAI, N=17; VAT: low VAI, N= 10 vs. low VAI =16) and *HSD11B1* (SAT: high VAI, N=17; VAT: low VAI, N= 12 vs. low VAI =17) (M). Low VAI; N =13 vs. high VAI, N= 18 unless otherwise stated.

## DISCUSSION

This study investigated steroid hormone metabolism and concentrations in women with severe obesity who were candidates for bariatric surgery. We found that in severe obesity, waist circumference alone or in conjunction with fasting serum triglycerides cannot identify individuals with adipose tissue dysfunction. Because we are looking at one extremity of the BMI value spectrum in this specific population, all individuals are abdominally obese according to the ATP-NCEP III criteria (51). We propose that other markers of adipose tissue dysfunction should be considered. VAI has been presented as an index of excess visceral adiposity associated with increased risk of cardiometabolic diseases (41). As reported in the expert consensus statement on abdominal obesity (50), adipose tissue dysfunction, ectopic fat deposition and cardiometabolic risk are intrinsically interconnected. In this study, we compared women having a significantly higher VAI to those with a low VAI. Our analyses revealed significant differences in plasma ApoB concentration and VAT adipocyte hypertrophy. This confirms previous literature showing that visceral adipose tissue hypertrophy is a marker of cardiometabolic status (7). Our results underline the notion that even in the context of severe obesity, the main features of adipose tissue dysfunction may still be relevant.

In addition, we found a significant difference in SAT and VAT *GPX3* gene expression between women with and without VAT excess. *GPX3* has been associated with insulin receptor downregulation in white adipose tissue in obese insulin-resistant mouse models and in humans with obesity (52). The same study demonstrated that silencing the *GPX3* gene decreases insulin receptor gene expression and its signaling cascade in 3T3-L1 adipocytes (52). Other data suggest that adipose *GPX3* is downregulated in cases of oxidative stress and inflammation (53). These results implicate *GPX3* in insulin resistance, reactive oxygen species (ROS) production and adipose tissue dysfunction. In our study, none of the 31 women were diabetic or taking hypoglycemics, suggesting that adipose *GPX3* gene expression is associated with local alterations in adipose tissue independent of these factors.

Interestingly, we found no significant difference in adipose tissue pericellular fibrosis between the two VAI groups. Pericellular fibrosis is thought to develop as a consequence of inadequate angiogenesis and unresolved low-grade inflammation, leading to pathological tissue remodeling and excessive extra-cellular matrix (ECM) deposition (54). Although this feature has been associated with adipose tissue dysfunction (55,56), we did not find any difference in pericellular fibrosis as a function of metabolic parameters in our study. Because we only included women with severe obesity, one possible explanation is that pericellular fibrosis is already high and no longer reflects intra-adipose dysfunction. Another possibility is that pericellular fibrosis, measured by the intensity of picrosirius red staining, reflects ECM remodeling rather than excess *per se*, and may therefore, not necessarily mirror adipose tissue dysfunction. Estrogens have been reported to stimulate ECM remodeling in breast cancer mouse models (57) and cultured fibroblasts (58). Here, we found that intra-adipose SAT estrone and estradiol amounts were associated with SAT pericellular fibrosis, consistent with the role of estrogens in ECM remodeling and adipose tissue fibrosis.

We conducted correlation analyses and mixed analyses to indirectly assess whether adipose tissue is actively involved in the uptake or release of active steroids or steroids precursors. We found no association between intra-adipose and plasma steroid concentrations, consistent with the notion that adipose tissue steroid hormones turnover is independent from plasma levels. This was already known for glucocorticoids (59) but in men, previous data showed positive correlations for estrone, testosterone and 5*α*-DHT (26). Differences in the techniques used (26) might explain the differences in the results. In addition, possible sex differences cannot be excluded. The presence and activity of most steroidogenic enzymes in adipose tissue has been confirmed and is reviewed in (60,61) and authors have suggested that adipocytes may also be able to participate in steroid *de novo* synthesis (62). Along with the ones examined here, other enzymes, such as 3*β*-hydroxysteroid dehydrogenase (3*β*HSD) converting dehydroepiandrosterone (DHEA) into A4, the 5*α*-reductase inactivating cortisol and transforming testosterone into 5*α*-DHT and the 17*β*HSD family may contribute to this lack of association (60,61).

It is generally accepted that in men, abdominal adiposity is inversely associated with plasma androgen concentrations, while the opposite is true for women (63). Our study found lower intra-adipose androgen concentrations, more specifically testosterone and androstenedione, in women with high VAI. In addition, we observed higher VAT AKR1C2 and aromatase activity in these women. Taken together, our data suggest the presence of increased indices of androgen aromatization (**Figure 2F-H** and **M**) and 5*α*-DHT catabolism (**Figure 2J**), which overall decrease the intra-adipose bioavailability of testosterone, androstenedione and 5*α*-DHT in women with increased visceral adiposity and adipose tissue dysfunction. Interestingly, despite a difference in terms of aromatase activity, our study did not identify differences in intra-adipose estrone or estradiol concentrations. This suggests that the feature associated with higher VAI is an overall decrease in intra-adipose androgen concentrations rather than an increase of estrogen concentrations.

Unfortunately, the number of studies that measured intra-adipose steroid concentration using gold-standard and reliable techniques using chromatography and mass spectrometry is scarce. This is mainly due to numerous challenges that adipose tissues and its matrix pose in extraction and quantification because other compounds present in the tissue may interfere with the steroid hormone signal during mass-spectrometry (42). One study on SAT androgen concentrations found that testosterone is increased and sequestered in adipocytes from men with obesity and insulin-resistant 3T3-L1 cells *in vitro* (64). However, it should be kept in mind that Di Nisio and colleagues (64) only measured SAT testosterone and compared groups of 6 patients, which could explain the differences between the two studies. Another study from our group conducted in women suggested that adipose tissues containing increased numbers of large, differentiated adipocytes likely display increased steroidogenic activity and in particular androgen catabolism (65). This was highlighted by increased expression and activity of AKR1C3, producing testosterone, and AKR1C2, the enzyme responsible for the reduction of 5*α*-DHT into its inactive metabolite 3α-androstenediol, and aromatase (65). Recently, we (34) and others (66) have demonstrated that adipose expression of *AKR1C2* and *AKR1C3*, positively associates with total and abdominal adiposity in women. Increasing androgen inactivation and lower testosterone concentrations were also reported by Marchand and collaborators (67), who demonstrated that plasma androstenedione in overweight pre-menopausal women correlated negatively with VAT area and, together with plasma testosterone, also with VAT hypertrophy (67). Taken together, our data and the current literature suggest that in women without PCOS, increased androgen catabolism is associated with visceral adiposity and adipose tissue dysfunction.

The intra-adipose balance between androgens and estrogens is modulated by aromatase *CYP19A1*, which converts testosterone and androstenedione into estradiol and estrone, respectively. Our data showed increased aromatase activity, mirrored by the estrogen-to-androgen ratio, and the specific increase of aromatase *CYP19A1* in VAT of women with visceral obesity and adipose tissue dysfunction. Aromatase is known to be significantly affected by total adipose tissue mass (68,69), probably due to the fact that its expression is increased with adipocyte differentiation (65). Indeed, both BMI and WC being indices of general and visceral adiposity, correlate positively with *CYP19A1* expression in pre-menopausal (70) but not post-menopausal women (19). Further supporting the effect of mass, recently Van de Velde and collaborators (71) reported that weight loss was accompanied by a decrease in plasma and urinary estrogen to androgen ratios suggesting a decrease in aromatase activity ratio in men who underwent bariatric surgery. Thus, the literature suggests a strong mass effect on aromatase expression and activity. However, our study revealed a specific increase in aromatase expression associated with VAT excess and dysfunction. In fact, although we did not find any differences in BMI or WC between women with high and low VAI, there was a significant difference in aromatase activity and VAT expression. In addition, VAT aromatase activity was associated with VAT hypertrophy even after WC adjustment while SAT activity was associated with dyslipidemia. Thus, in our study, adipose tissue mass alone could not completely explain the increased aromatase activity and expression observed.

Androgen concentrations and metabolic impairments are particular relevant for women with PCOS, as these patients are often characterized by abdominal obesity (63) and features of adipose tissue dysfunction, such as enlarged adipocytes (72,73). Current literature indicated increased AKR1C2, 3*β*HSD (74) and AKR1C3 (74,75), in SAT of PCOS women compared to those without PCOS. On the other hand, aromatase expression seems lower in women with PCOS (74). In addition, it seems that *in vitro* differentiated primary preadipocyte from PCOS patients produce significantly more testosterone than the non-PCOS controls (75). In order to tackle excess androgen production in PCOS, patient are sometimes prescribed flutamide, an androgen receptor inhibitor. Interestingly, flutamide not only can lower testosterone action in target tissues but may also help improve the cardiometabolic alterations (76) and features of adipose tissue dysfunction (77) observed in PCOS. Our study was not designed to compare PCOS to non-PCOS women with severe obesity. However, the adipose tissue features that we observed in women with a high VAI are not fully consistent with those expected in women with PCOS.

Our study has a number of strengths. First, we were able to measure intra-adipose steroid hormones using novel and highly sensitive techniques (42). Similarly, AKR1C2 activity was measured using a fluorogenic probe which has demonstrated specificity for the enzyme in question (46), and has never been used on adipose tissue homogenates. Both techniques are novel and contribute to the technological advance in the characterization of adipose tissue steroid concentrations and inferred AKR1C2 activity. In addition, thanks to our study design, we were able to measure and compare paired intra-adipose and plasma steroids. Our study also has limitations. Although in this study we did not measure matrix effect by LC-ESI-MS/MS, methods upon which our study is based showed that the current protocol reduces ion interference without eliminating it. We included only 31 women and the number of men was too small to perform further analysis, preventing us to accurately compare steroidogenic homeostasis in men and women with VAT excess and adipose tissue dysfunction. In addition, as anthropometric data were collected retrospectively, we have no information on adipose tissue distribution performed using gold-standard methods (1). To counteract this, we used the VAI, which has been described as a reliable estimator of VAT area, volume and adipose tissue dysfunction (41). In addition, we extensively characterized adipose tissue dysfunction markers including measurements of adipocyte hypertrophy, pericellular fibrosis and the expression of *DGAT2*, involved in triglyceride biosynthesis, and *GPX3*, implicated in intracellular ROS balance and adipose insulin resistance.

In conclusion, we found that VAT aromatase activity is positively associated with VAT adipocyte hypertrophy and negatively with plasma HDL-cholesterol, while SAT aromatase activity predicted dyslipidemia in women even after adjustment for WC, age and anovulant intake. We additionally compared women with high and low VAI and found that VAT excess is characterized by features such as adipose tissue dysfunction, decreased presence of active androgens and increased aromatase activity and expression. Overall, our data question the assumed deteriorating effect of androgens on adipose tissue function in women and suggest that the balance between androgens and estrogens might be relevant in women without PCOS.

## AKNOWLEDGMENTS

The Authors acknowledge the invaluable collaboration of the surgery team, bariatric surgeons and biobank staff of the IUCPQ.

## DATA AVAILABILITY STATEMENT

Datasets analyzed for the purposes of the current study are available from the corresponding author on reasonable request. However, to preserve patient confidentiality, restrictions may apply.

