## Supplemental Table S1 for "Increased adipose tissue indices of androgen catabolism and aromatization in women with metabolic dysfunction"

**SUPPLEMENTARY DATA**

**Table S1: Lower limit of quantification for adipose and plasma steroids.**

| **Sample** | **Steroid** | **Lower limit of quantification** |
| --- | --- | --- |
| Adipose tissue | Estrone | 15 pg |
|  | Estradiol | 25 pg |
|  | Testosterone | 25 pg |
|  | 5α-dihydrotestosterone | 25 pg |
|  | Androstenedione | 100 pg |
|  | Dehydroepiandrosterone | 100 pg |
|  | Cortisone | 75 pg |
|  | Cortisol | 100 pg |
| Plasma | Estrone | 0.0125 ng/mL |
|  | Estradiol | 0.0125 ng/mL |
|  | Cortisone | 0.0125 ng/mL |
|  | Cortisol | 0.0125 ng/mL |
|  | 5α-dihydrotestosterone | 0.025 ng/mL |
|  | Testosterone | 0.05 ng/mL |
|  | Androstenedione | 0.05 ng/mL |
