## Supplemental Table S2 for "Increased adipose tissue indices of androgen catabolism and aromatization in women with metabolic dysfunction"

**Table S2: List of primers used**. Primers were designed using NCBI primer design tool and validated by PCR before use.

| **Gene** | **GenBank reference** | **Size (base pairs)** | **Sequence** (5‘ 🡪 3‘// 3‘🡪5‘) |
| --- | --- | --- | --- |
| GAPDH | NM_002046 | 194 | GGCTCTCCAGAACATCATCCCT // ACGCCTGCTTCACCACCTTCTT |
| AKR1C2 | [NM_001354.6](https://www.ncbi.nlm.nih.gov/entrez/viewer.fcgi?db=nucleotide&id=1890346376) | 341 | GCTCTTATAGCCTGTGAGGGAG // GACCAACTCTGGTCGATGGG |
| AKR1C3 | NM_003739.6 | 150 | ACAGGGAATGGATTCCAAACACCA // TGGCGGAACCCAGCTTCTAT |
| CYP19A1 | NM_031226.3 | 333 | TGAATCGGGCTATGTGGACG // AGGTCACCACGTTTCTCTGC |
| HSD11B1 | NM_005525.4 | 269 | CATTGCTGGCACCATGGAAG// GATAAGCCACTTTCCCAGCCA |
| GPX3 | NM_002084.5 | 171 | GTACGGAGCCCTCACCATTG // AAGCCCAGAATGACCAGACC |
| PLOD2 | [NM_000935.3](https://www.ncbi.nlm.nih.gov/entrez/viewer.fcgi?db=nucleotide&id=169791021) | 316 | TTACGGCAAATGGTCTGGGG // GCAACCACCTCCCTGAAAGT |
| DGAT2 | NM_032564.5 | 178 | ACACACCCAAGAAAGGTGGCA // AAGTTGCAGAAGGCACCCAG |
