## Supplemental Table S2 for "Increased adipose tissue indices of androgen catabolism and aromatization in women with metabolic dysfunction"

**Table S3: Comprehensive summary of men and women included in this study.** Data are represented as mean ± standard deviation unless otherwise stated.

|  | **Women (N=31)** | **Men (N=9)** | **All (N=40)** |
| --- | --- | --- | --- |
| Age (years) | 39.1 ± 7.0 | 40.9 ± 9.7 | 39.5 ± 7.6 |
| BMI (kg/m2) | 50.6 ± 5.7 | 48.5 ± 8.5 | 50.1 ± 6.4 |
| WC (cm) | 131.7± 11.2 | 140.7 ± 15.2 | 133.6 ± 12.4 |
| Hip circumference (cm) | 148.2 ± 12.7 | 134.6 ± 12.1 | 145.7 ± 13.6 |
| Waist-to-hip ratio | 0.89 ± 0.07 | 1.04 ± 0.04 | 0.92 ± 0.08 |
| Neck circumference (cm) | 41.9 ± 3.2 | 43.2 ± 3.2 | 43.2 ± 4.0 |
| SAT adipocyte cell diameter (um) | 91.8 ± 12.3 | 88.3 ±8.7 | 90.9 ± 11.4 |
| VAT adipocyte cell diameter (um) | 83.5 ± 14.14 | 90.8 ± 7.1 | 85.3 ± 13.0 |
| SAT pericellular fibrose (%) | 6.0 ± 1.7 | 6.1 ± 1.2 | 6.1 ± 1.6 |
| VAT pericellular fibrose (%) [Md ±IQR] | 3.9 ± 1.7 | 3.6 ± 2.6 | 3.9 ± 1.9 |
| Fasting glucose (mM) | 5.8 ± 0.6 | 5.7 ± 0.64 | 5.8 ± 0.6 |
| HbA1c (%)[Md ±IQR] | 5.7 ± 0.4 | 5.3 ± 0.7 | 5.6 ± 0.4 |
| Fasting triglycerides (mM) [Md ±IQR] | 1.55 ± 0.94 | 1.48 ± 0.76 | 1.46 ± 0.92 |
| Total cholesterol (mM) | 4.66 ± 0.63 | 5.12 ± 0.86 | 4.75 ± 0.70 |
| HDL-c (mM) | 1.20 ± 0.31 | 1.03 ± 0.09 | 1.17 ±0.29 |
| LDL-c (mM) | 2.75 ± 0.52 | 3.56 ± 0.44 | 2.91 ± 0.59 |
| Hypoglycémiants (Y:N) | (0:31) | (0:9) | (0:40) |
| Hypolipidemians (Y:N) | (0:31) | (0:9) | (0:40) |
| Corticosteroids (Y:N) | (0:31) | (0:9) | (0:40) |
| CAR AUCi [Mdn ± IQR] | 92.4 ± 129.5 | 89.8 ± 14.8 | 92.4 ± 146.7 |
| S1 mean (nM) [Mdn ± IQR] | 11.5 ± 5.9 | 13.3 ± 9.5 | 11.6 ± 6.3 |
| S2 mean (nM) [Mdn ± IQR] | 14.3 ± 8.4 | 15.3 ± 8.1 | 14.3 ± 7.9 |
| S3 mean (nM) | 16.0 ± 8.3 | 18.0 ± 6.5 | 16.4 ± 7.9 |
| Plasma cortisol (nM) | 217.7 ± 92.1 | 239.4 ± 80.3 | 222.1 ± 89.2 |
| Plasma cortisone (nM) | 54.5 ± 19.3 | 70.5 ± 15.5 | 57.8 ± 19.5 |
| Plasma testosterone (pM) | 751.4 ± 275.6 | 9 999.0 ± 2646.4 | 2750.9 ± 4039.4 |
| Plasma dihydrotestosterone (pM) | 209.0 ± 82.2 | 844.2 ± 258.6 | 339.3 ± 291.9 |
| Plasma estrone (pM) | 718.2 ± 471.7 | 264.1 ± 82.7 | 625.1 ± 459.8 |
| Plasma estradiol (pM) | 432.5 ± 239.1 | 236.0 ± 72.5 | 391.1 ± 228.9 |
| SAT cortisol (pmol/kg) | 65591.0 ± 70957.0 | 107672.0 ± 68430.1 | 72166.0 ± 71190.1 |
| VAT cortisol (pmol/kg) | 64065.0 ± 57712.8 | 83312.0 ± 23894.6 | 67674.0 ± 53258.3 |
| SAT cortisone (pmol/kg) | 23132.0 ± 23979.8 | 44028.0 ± 27364.0F | 28663.0 ± 26216.3 |
| VAT cortisone (pmol/kg) | 23131.6 ± 30748.7 | 30922.0 ± 19610.5 | 29332.0 ± 28599.8 |
| SAT androstenedione (pmol/kg) [Md±IQR] | 26592.0 ± 65735.8 | 56255.0 ± 26805.5 | 33155.0 ± 58673.1 |
| VAT androstenedione (pmol/kg) | 57212.0 ± 57337.6 | 66787.0 ± 36393.3 | 59228.0 ± 53319.8 |
| SAT testosterone (pmol/kg) | 2545.0 ± 3317.4 | 10816.3 ± 9412.7 | 3518.1 ± 5004.2 |
| VAT testosterone (pmol/kg) | 2045.0 ± 2068.1 | 8567.0 ± 696.1 | 2453.4 ± 2566.9 |
| SAT dihydrotestosterone (pmol/kg) [Md ±IQR] | 630.9 ± 450.8 | 1722.8 ± 981.1 | 830.3 ± 1126.4 |
| VAT dihydrotestosterone (pmol/kg) [Md±IQR] | 473.3 ± 547.7 | 1967.1 ± 1152.9 | 1137.1 ± 1542.1 |
| SAT estrone (pmol/kg) | 5375.0 ± 3922.7 | 5317.3 ± 3372.5 | 5362.0 ± 3764.3 |
| VAT estrone (pmol/kg) | 5382.0 ± 3517.5 | 6160.0 ± 3533.2 | 5525.3 ± 3485.7 |
| SAT estradiol (pmol/kg) | 3133.9 ± 2360.4 | 3591.8 ± 1879.8 | 3442.4 ± 2240.4 |
| VAT estradiol (pmol/kg) | 3270.0 ± 2535.4 | 3240.3 ± 1755.7 | 3264.0 ± 2361.3 |

Abbreviations: WC= waist circumference; SAT= subcutaneous adipose tissue, VAT = visceral adipose tissue, HDL-c = high-density lipoprotein cholesterol, LDL-c = low-density lipoprotein cholesterol, CAR= Cortisol awakening response, AUCi= incremental area under the curve Mdn = median, IQR = inter-quartile range
